# Development of the mandibular curve of Spee and maxillary compensating curve: A finite element model

**DOI:** 10.1101/723429

**Authors:** Steven D. Marshall, Karen Kruger, Robert G. Franciscus, Thomas E. Southard

## Abstract

The curved planes of the human dentition seen in the sagittal view, the mandibular curve of Spee and the maxillary compensating curve, have clinical importance to modern dentistry and potential relevance to the craniofacial evolution of hominins. However, the mechanism providing the formation of these curved planes is poorly understood. To explore this further, we use a simplified finite element model, consisting of maxillary and mandibular “blocks”, developed to simulate tooth eruption, and forces opposing eruption, during simplified masticatory function. We test our hypothesis that curved occlusal planes develop from interplay between tooth eruption, occlusal load, and mandibular movement.

Our results indicate that our simulation of rhythmic chewing movement, tooth eruption, and tooth eruption inhibition, applied concurrently, results in a transformation of the contacting maxillary and mandibular block surfaces from flat to curved. The depth of the curvature appears to be dependent on the radius length of the rotating (chewing) movement of the mandibular block. Our results suggest mandibular function and maxillo-mandibular spatial relationship may contribute to the development of human occlusal curvature.

## Introduction

Viewed in the sagittal plane, the concave occlusal curvature of the mandibular dentition is a naturally occurring phenomenon seen in modern humans and fossil hominins [1,2], Commonly described as the *curve of Spee* [3,4], this occlusal curvature (Fig 1) is proposed to have functional significance during mastication [5-8] and is considered an important element of diagnosis and treatment planning in dentistry and orthodontics. [7-18]

**Fig 1.**
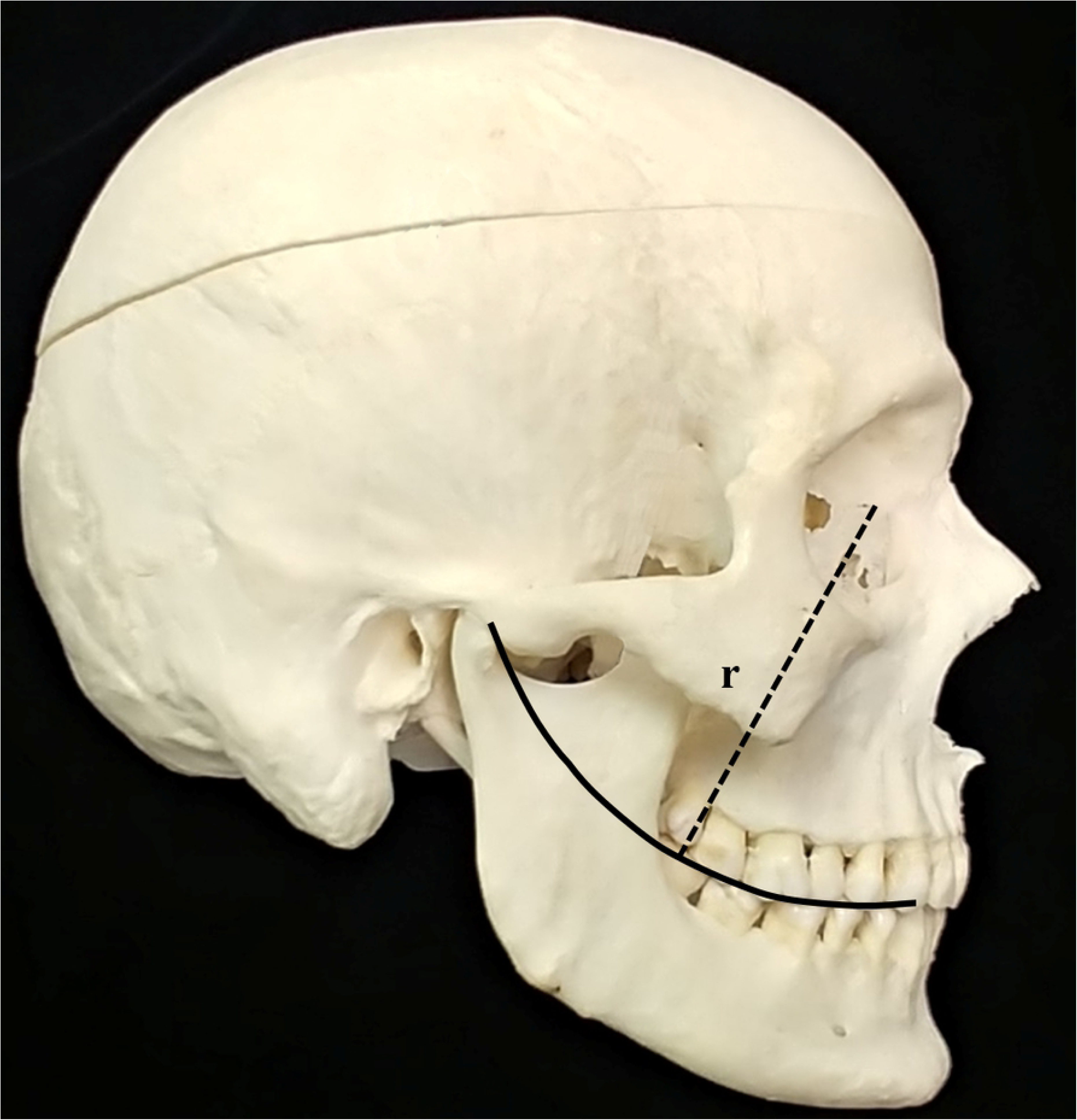
Human skull in norma lateralis depicting the antero-posterior curvature of the maxillary and mandibular dentition. Dotted line depicts curve of Spee radius of length “r”.

Despite its clinical importance in extant humans, and its potential relevance to fossil hominin craniofacial evolution, the process by which the concave mandibular curve of Spee develops is not completely understood. Many investigations have analyzed variables thought to be associated with the curve’s development, including the timing of permanent tooth eruption and vertical alveolar development [19,20], variation in craniofacial morphology and various factors associated with malocclusion [2,14,17,20-22], but the underlying mechanisms of the curve’s formation and maintenance remain unknown. In a similar fashion, while intuitively logical, the process by which the corresponding maxillary convex occlusal curvature (i.e. the compensating curve, Fig 1) develops is, nonetheless, also not understood. Unanswered are the following questions: why should a concave curve develop in the mandibular dental arch, why should a convex curve develop in the maxillary arch, why should these two curves develop simultaneously, and why should they persist throughout life?

The development of these curves, in part, must be related to tooth eruption. Post-emergent tooth eruption (i.e. tooth eruption after a tooth has emerged through gingival tissue) has been divided into four stages: pre-functional spurt, juvenile equilibrium, adolescent spurt, and adult equilibrium, with the last three stages designating eruptive tooth movement after the tooth has reached the occlusal plane. [27] As teeth erupt, they appear to erupt until they find their position along these curves. In addition, the development of these curves must be related to occlusal forces acting against the teeth. For instance, when a maxillary or mandibular molar is unopposed by another tooth, the molar will continue to erupt (i.e., super-eruption), irrespective of the curves formed from the remaining teeth. In other words, occlusal loading must be involved in development of the curves. Lastly, it seems likely that the development of these curves must be related to mandibular movement, constrained within a pathway determined by muscular and ligamentous attachments, either during function or during parafunction, the latter including bruxism and other activities not related to eating, drinking, or talking.

Considering the application of occlusal forces to the teeth during functional and parafunctional tooth contact, and the equilibrium theory of tooth position [28], it is conceivable that the development and maintenance of the curve of Spee and compensating curve, results from the interplay between tooth eruption, occlusal loading, and mandibular movement during function.

The purpose of this investigation is to test the hypothesis, using a simplified finite element model (FE model), that mandibular curve of Spee and maxillary compensatory curve develop from a predictable interplay between tooth eruption, occlusal load, and mandibular movement.

## Materials and methods

The finite element method uses discretization to compute approximations to space- and time-dependent problems. It has been widely-used in studying the functional morphology of the human masticatory system.[29]

We based our FE model upon the following assumptions:

- The curve of Spee, as classically described by Ferdinand Graf von Spee (Fig 1), exists as a *concave* curve in the mandibular dental arcade, with an average radius of approximately 100 mm [30]. A corresponding *convex* curve (compensating curve) exists in the maxillary dental arcade.
- Both curves result from an interplay between tooth eruption, inhibition of tooth eruption by axial occlusal loading, and mandibular movement during function.
- Maxillary and mandibular teeth erupt toward the occlusal plane at a continuous rate (E_C_).
- Occlusal loading occurs during mastication as the mandibular dental arcade rotates around a center of rotation (C_ROT_) that lies above the maxillary dentition. The position of C_ROT_ results from the spatial relationship of the condyles to the mandibular dental arcade and the complex interaction between condylar head translation, condylar head rotation, muscle tension, and ligament restriction.
- The inhibition of tooth eruption (a relative apical tooth movement) is a product of the magnitude of occlusal loading at each point of occlusal contact in a direction normal to the occlusal surface (F_n_) integrated throughout the time this force is applied (T).
- Total tooth eruption (E_TOTAL_) at any point along the maxillary and mandibular dental arch is calculated as the difference between tooth eruption toward the occlusal surface and inhibition of tooth eruption or,

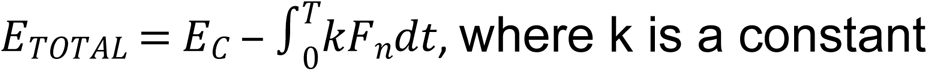
- Our FE model is restricted along the 2 dimensions of the sagittal plane

Our FE model was created using ABAQUS v6.9.1 and consisted of two blocks of elements, the upper block representing the maxillary dentition and the lower block representing the mandibular dentition. The block’s left side represents posterior teeth and the block’s right side represents anterior teeth.. Each block was composed of a single layer of brick continuum elements (σ=82.5 GPA, v=0.33) [31] with 40 elements horizontal, 20 elements vertical, and 1 element deep (Fig 2). That is, 1600 elements are found within each maxillary and mandibular block resulting in block dimensions of 20 mm high by 40 mm wide (40 mm is an approximation for posterior-anterior human dental arch length). The maxillary block was held fixed while the mandibular block was rotated about C_ROT_, simulating simplified human chewing motion in 2 dimensions from the sagittal view. Two centers of mandibular block rotation were simulated: one at 100 mm (an approximation of the human curve of Spee), and one at 400 mm (a value chosen to evaluate the behavior of our FE model with increasing C_ROT_), above the superior surface of the mandibular block (i.e. occlusal surface of the mandibular dentition). The starting form of the maxillary and mandibular blocks was rectangular in shape (i.e. 40w × 20h) before being subjected to motion, simulated tooth eruption and masticatory load. One rotation cycle is described as follows: The cycle begins with the maxillary and mandibular blocks in contact such that the posterior surface of the mandibular block approximately 3 mm posterior to the posterior surface of the maxillary block (Fig 2C). The cycle proceeds with the rotation of the mandibular block, along an arc (i.e. the rotation arc) dictated by the length of the radius between C_ROT_ and the midpoint of the occlusal surface of the mandibular block, to a point where the anterior surface of the mandibular block is approximately 3 mm anterior to the anterior surface of the maxillary block (Fig 2D). Each rotation cycle, the blocks are moved toward each other (maxillary block moves inferiorly, mandibular block moves superiorly) a distance of 0.01 mm (our proxy for tooth eruption). Similarly, during each rotation cycle, block material was removed at the occlusal surface (our proxy for tooth eruption inhibition) based on a calculation using the Archard-Lancaster relationship for wear. [32] An ABAQUS UMESHMOTION subroutine was created which allowed for this simulated eruption inhibition on both plates as sliding occurred while they were in contact. Following each rotation cycle, the subroutine computed the new occlusal surface topography (depth for each surface node) and the model was re-meshed to simulate this modified geometry prior to entry into the subsequent cycle. In our study, the blocks were modeled as identical material (which would be the case for tooth composition in both arches) and the subroutine had to be adapted to permit maxillary and mandibular blocks to undergo eruption-inhibition at equal rates (for similar values of normal load and time of load application). The large shape change experienced by the “occlusal” surface of the maxillary and mandibular blocks required intensive remeshing of both the surface and underlying nodes to preserve element quality throughout the simulation. The simulation was run until a steady-state curve was reached. The length of the mandibular block rotation arc was chosen to allow a reasonable time to reach steady-state.

**Fig 2.**
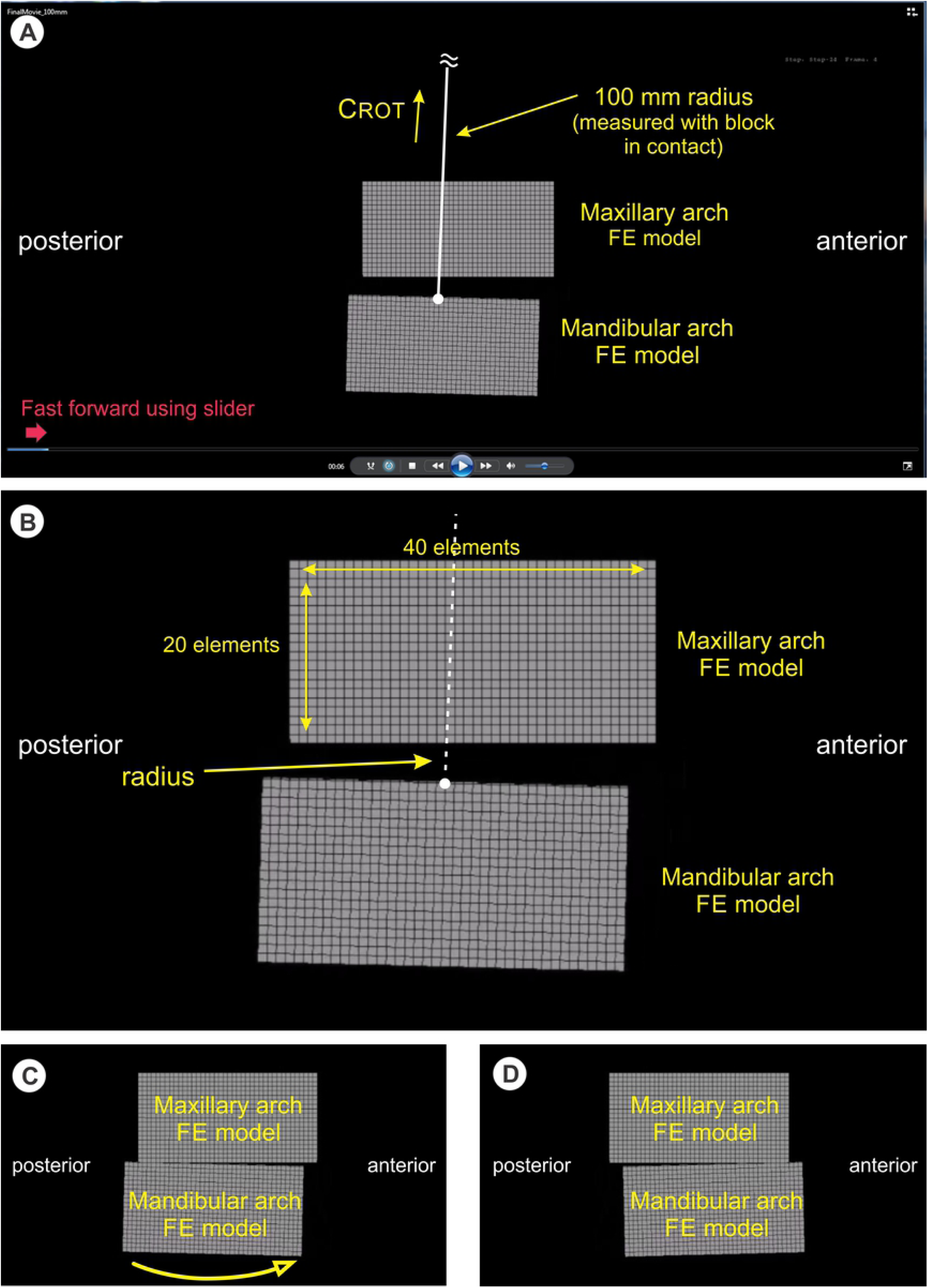
Diagram explaining visualization of FE model video and definition of terms. (A) The mandibular arch FE model has passed through one rotation arc (left to right), has disengaged with the maxillary arch FE model, and is now returning to the left to engage the maxillary arch FE model for a second cycle. Using the mouse cursor (red arrow) to engage and move slider bar allows fast-forwarding of the movie to quickly see the block shape effect of the cycles with time. CRoT for the m andibu lar arch FE model lies 100 mm above the center of the superior surface of the mandibular block when it is aligned vertically, directly underneath and in contact with, the maxillary arch FE model. (B) Closer view of Figure 2A showing the dimensions and arrangement of the elements in the FE models. Each block is 40 elements in width, 20 elements in length, and 1 element in depth. (C) Position of mandibular arch FE model before sliding across the inferior surface of the maxillary arch FE model along the rotation arc dictated by the radius to CR oT• (D) Position of the mandibular arch FE model after sliding across the inferior surface of the maxillary arch FE model, completing one rotation arc cycle.

## Results

Simulations of both models (C_ROT_ = 100m and C_ROT_ = 400 mm, in mp4 movie format) may be observed at the web sites shown in Figure 3. An application that allows viewing of mp4 files is required (e.g. Windows Media Player®, QuickTime®, iTunes®). To move quickly through the simulation, advance the time slider at the bottom from left to right (see Fig 2A).

**Fig 3.**
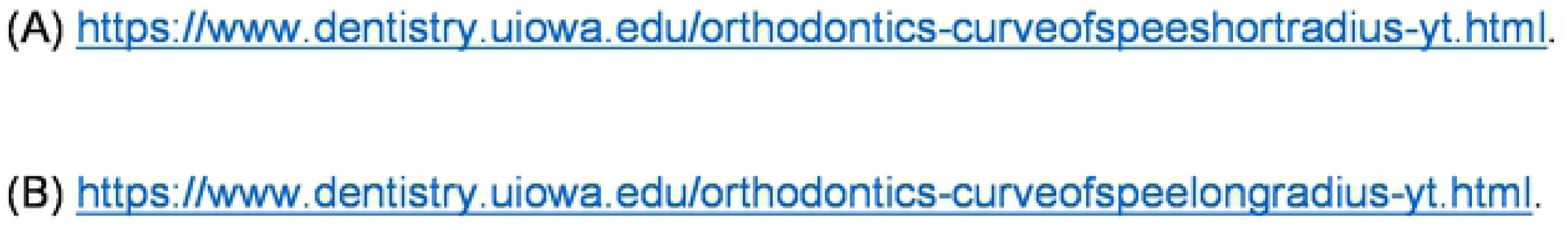
World Wide Web address of the FEM simulation of human masticatory movements seen in the sagittal plane. (A) https://www.dentistrv.uiowa.edu/orthodontics-curveofspeeshortradius-yt.html. Radius of mandibular chewing rotation = 100 mm. (B)https://www.dentistry.uiowa.edu/orthodontics-curveofspeelongradius-yt.htmI. Radius of mandibular chewing rotation = 400 mm.

As observed in the simulations, repetitive rotation cycles cause the surfaces of the maxillary and mandibular blocks (representing the maxillary and mandibular dentitions) to undergo a notable change that eventually reaches equilibrium. Two-dimensional rotation of the mandibular block against the stationary maxillary block during simulated tooth eruption interacting with simulated tooth eruption inhibition, resulted in curvature of the contacting block surfaces (i.e. occlusal surfaces) (Fig 3). Beginning with a flat occlusal surface, as the mandibular dentition undergoes a chewing rotation about C_ROT_, a concave occlusal surface (curve of Spee) develops in the mandible, and a compensating curve develops in the maxilla. When the rotation radius of the mandibular block is smaller, the resulting maxillary and mandibular block curves are deeper (Fig 3A). When the rotation radius of the mandibular block is larger, the resulting upper and lower block curves are shallower (Fig 3B).

## Discussion

Using a FE model we sought to simulate the interplay of forces during mastication: tooth eruption modeled by maxillary FE model brick continuum elements being brought towards mandibular FE model brick continuum elements with each cycle; and tooth eruption inhibition modeled as a function of axial loading during a simplified rotational masticatory movement. The spatial relationships of this simplified FE model approximated human maxillo-mandibular spatial relationship as seen from the sagittal view, with the mandibular rotation radius = 100 mm, and the dental arcade length from the sagittal view = 40 mm. The rotation arc of approximately 3 mm is large compared to that shown for humans (≈ 0.5 mm) in the sagittal plane [24,25], and was chosen to allow a reasonable length of time for our simulation to reach steady state.

The principle finding of our pilot study is that the surfaces of the maxillary and mandibular FE model blocks, in contact during simulation of human chewing in two dimensions during which the forces of tooth eruption and tooth loading are also simulated, transform from their original flat shape to a curved shape in the steady-state. We find steady-state curve shape is dependent on the rotation arc radius of the mandibular FE model block, with a smaller rotation arc radius producing with a deeper curve. These findings have implications for explaining variation in the development and maintenance of the mandibular curve of Spee and maxillary compensating curve in humans. A depiction of the variation in curves generated by our analysis, displayed as maxillary and mandibular dentitions, is shown in Figure 4.

**Fig 4.**
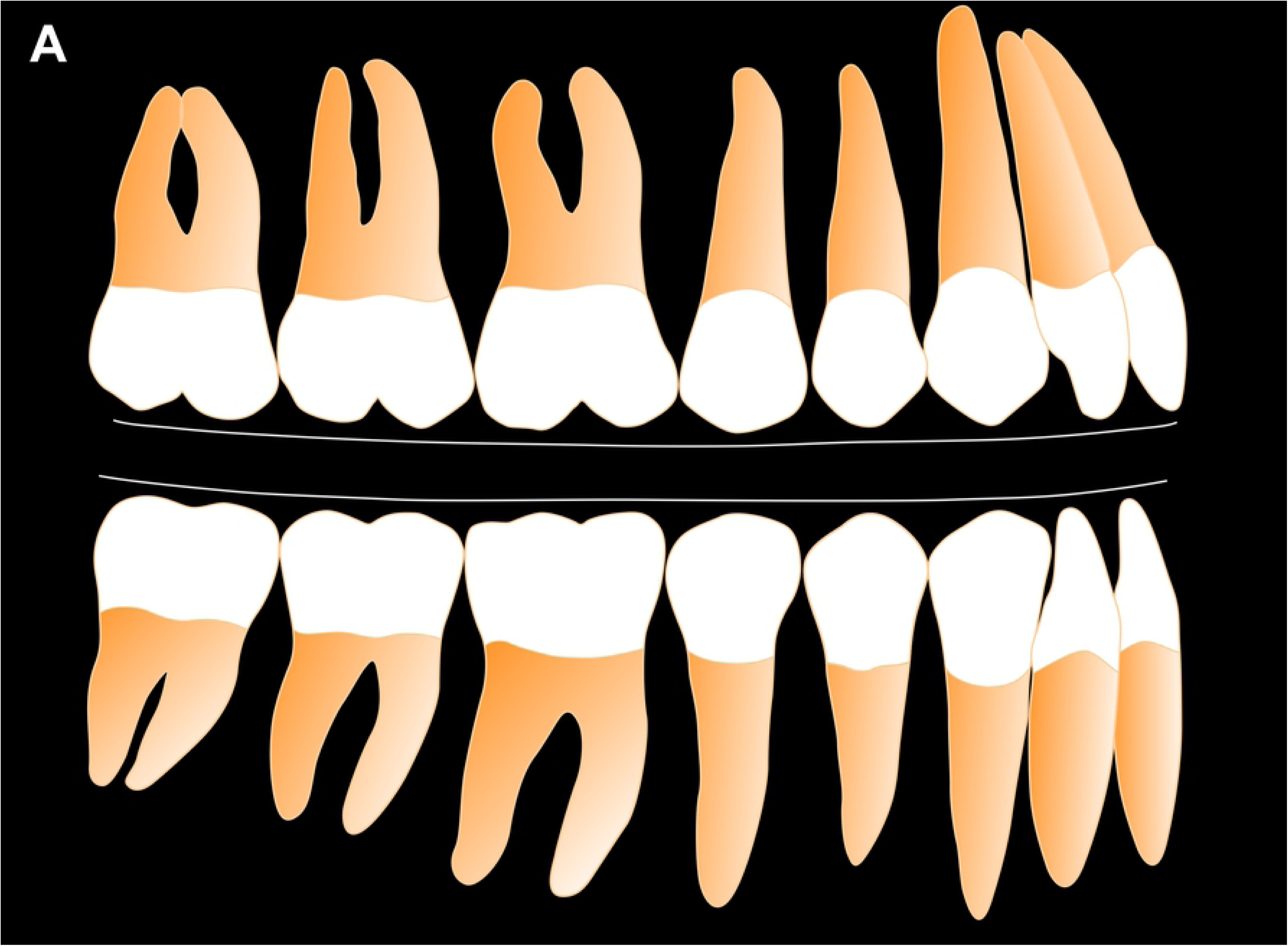
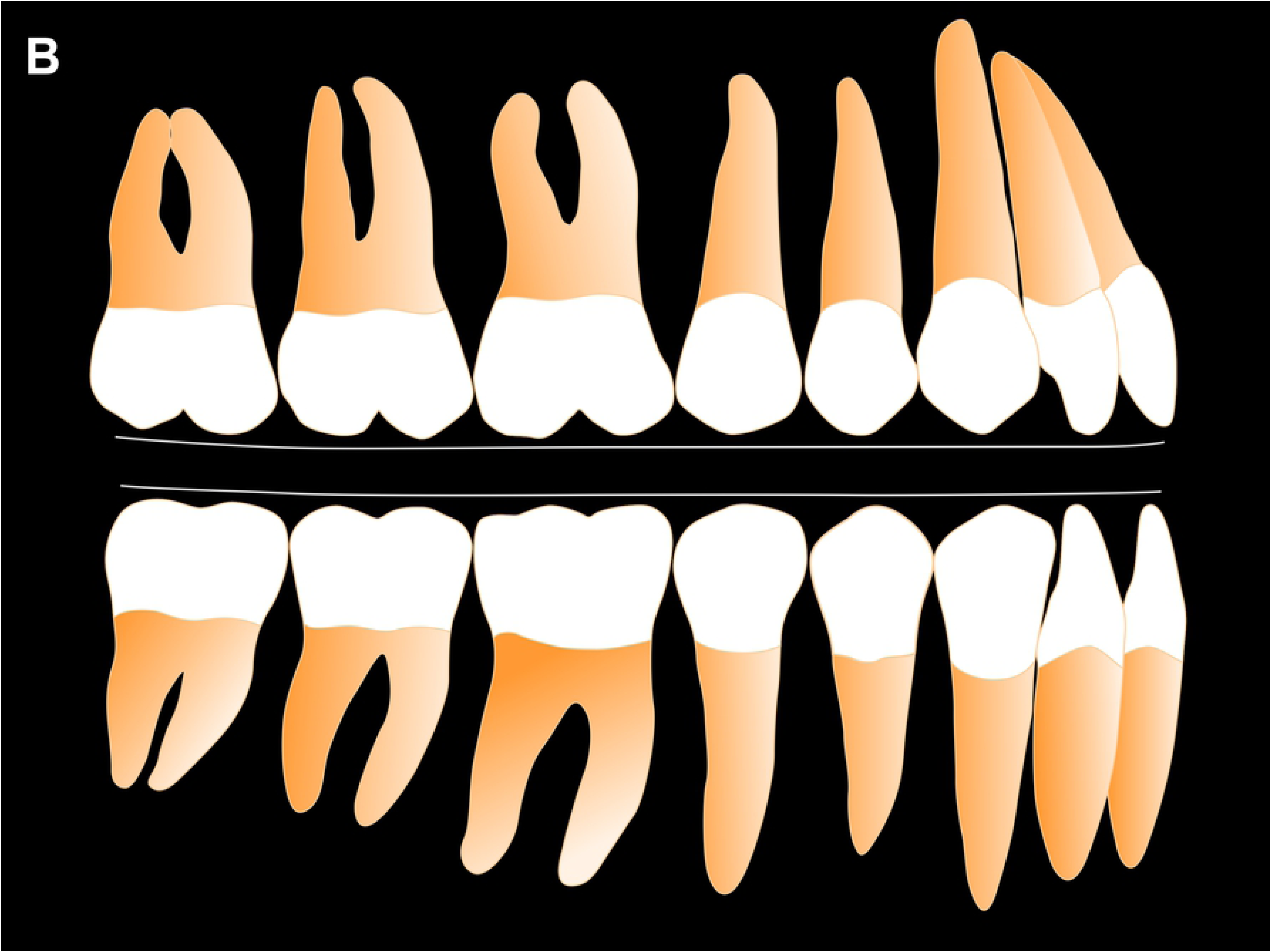
Depiction of occlusal curvature (mandibular curve of Spee, maxillary compensating curve) developed by FEM simulation seen in Figs 3A and 3B. **(A)** C_RoT_ = 100 mm. (B) C_RoT_ = 400 mm.

Our model is based, in part, on the equilibrium theory of tooth position, the assumption that eruptive forces continue throughout life and are opposed by forces generated during dynamic contact and release of mandibular agonist teeth with their maxillary antagonists. [26,27] Although, the direction, magnitude and duration of force necessary to inhibit eruptive tooth movement is not fully understood [27], we assume that the loading of the teeth during the rhythmic mandibular movements of mastication, and daytime and nocturnal bruxism, results in complex loading of the maxillary and mandibular teeth, including forces directed apically along the long axis of individual teeth (i.e. axial loading). [23,24,25,40-42] There exists ample evidence that tooth position equilibrium is disrupted when axial loading forces are greatly diminished. Numerous studies of humans and fossil hominids have demonstrated the super-eruption of teeth when occlusal antagonist teeth are lost. [33-39]

The influence of the spatial relationship of the mandibular condyles to the dentition is not well understood. Studies have suggested the horizontal distance between the condyle and the dentition is inversely related to the depth of the curve of Spee. [2,5,6,11,14,17] One theoretical explanation for these observations suggests a shorter horizontal distance between the condyle and the dentition necessitates a deeper curve of Spee for greater mechanical advantage during chewing. [5,6] Our FEM simulation did not test this directly. A more complex FE model is required to address this question.

## Conclusions

A finite element model (FEM) was developed to simulate tooth eruption, and forces opposing eruption, during simplified masticatory function. The maxillary arch FE model was held stationary, and the mandibular arch FE model was moved against the maxillary block to simulate two-dimensional chewing motion. The results of our experiment show:

1. Two-dimensional simulation of rhythmic chewing movement, tooth eruption, and tooth eruption inhibition applied concurrently, resulted in a transformation of the contacting block surfaces from flat to curved.
2. A shorter radius for the rotation arc of the mandibular dental arch results in more deeply curved surfaces of the maxillary and mandibular dental arches. A larger radius for the rotation arch of the mandibular dental arch results in shallower curved surfaces of the maxillary and mandibular dental arches. This finding may explain, in part, human mandibular curve of Spee and maxillary compensating curve variation.

## Authors’ contribution

S.D.M., K.K., R.G.F., and T.E.S. have contributed equally to this work. S.D.M. contributed conceptualization, supervision, validation, visualization, original draft preparation. K.K. contributed data curation, formal analysis, investigation, software, validation, review & editing. R.G.F. contributed conceptualization, validation, visualization, review & editing. T.E.S. contributed conceptualization, methodology, resources, supervision, validation, visualization, review & editing.

